# Genomic background sequences systematically outperform synthetic ones in de novo motif discovery for ChIP-seq data

**DOI:** 10.1101/2023.12.30.573742

**Authors:** Vladimir V. Raditsa, Anton V. Tsukanov, Anton G. Bogomolov, Victor G. Levitsky

**Author notes:** Victor G. Levitsky, Department of System Biology, Institute of Cytology and Genetics, Novosibirsk, 630090, Russia.

## Abstract

Efficient *de novo* motif discovery from the results of wide-genome mapping of transcription factor binding sites (ChIP-seq) is dependent on the choice of background nucleotide sequences. The foreground sequences (peaks) represent not only specific motifs of target transcription factors, but also the motifs overrepresented throughout the genome, such as simple sequence repeats. We performed a massive comparison of the ‘synthetic’ and ‘genomic’ approaches to generate background sequences for *de novo* motif discovery. The ‘synthetic’ approach shuffled nucleotides in peaks, while in the ‘genomic’ approach randomly selected sequences from the reference genome or only from gene promoters according to the fraction of A/T nucleotides in each sequence. We compiled the benchmark collections of ChIP-seq datasets for mammalian and Arabidopsis, and performed *de novo* motif discovery. We showed that the genomic approach has both more robust detection of the known motifs of target transcription factors and more stringent exclusion of the simple sequence repeats as possible non-specific motifs. The advantage of the genomic approach over the synthetic one was greater in plants compared to mammals. We developed the AntiNoise web service (https://denovosea.icgbio.ru/antinoise/) which implements a genomic approach to extract genomic background sequences for twelve eukaryotic genomes.

## INTRODUCTION

Transcription factors (TFs) are proteins that control gene transcription by sequence-specific DNA binding. Chromatin immunoprecipitation (ChIP)-based high throughput technique ChIP-seq allows genome-scale mapping of TF binding sites (TFBS) (1,2) (Nakato and Shirahige, 2017; Lloyd and Bao, 2019). The primary processing of raw ChIP-seq data is called peak calling. It produces thousands of genomic loci or peaks enriched by protein binding (3,4 Tran and Huang, 2014; Thomas et al., 2017). Each peak is usually at least several hundred base pairs long; it usually has a clear point of a number of maximum number of reads (‘summit’), presumably correlating with positioning of motifs of target TFs (5,6 Pepke et al., 2009; Kulakovskiy and Makeev, 2013). The term target TF refers to the TF studied in the ChIP-seq experiment. The most important task of secondary processing is to detect in the peaks specific motifs that are responsible for biological functions of the target TF. A specific motif represents the short, recurring pattern that is thought to have a biological function of sequence-specific binding sites for TF (7,6 D’haeseleer, 2006; Kulakovskiy and Makeev, 2013). Conventionally, a motif varies in length from 6 to 20 bp (8-10 Spitz and Furlong 2012; Zambelli et al., 2013; Vorontsov et al., 2024). The motif enrichment analysis, and above all, *de novo* motif discovery, is the required step to map exact positions of potential TFBS in peaks (3,11 Tran and Huang, 2014; Liu et al., 2018). An aim of motif discovery consists in exact definition of all parameters of a motif model.

Over the last few years, thousands of uniformly processed ChIP-seq datasets for hundreds of target TFs for major model species, including mammals, plants, insects, worms, and fungi have been collected in several focused databases (Cistrome DB, ReMap and GTRD, 12-14 Taing et al., 2019; Hammal et al., 2022; Kolmykov et al., 2021). Also, the massive application of *in vitro* technologies such as PBM (Protein Binding Microarray) and HT-SELEX (High-Throughput *in vitro* Selection) detected specific sequence motifs for several hundreds of TFs (15-18 Stormo and Zhao, 2010; Jolma and Taipale, 2011; Jolma et al., 2013; Franco-Zorrilla et al., 2013). As a result, the public databases JASPAR (19 Rauluseviciute et al., 2024), Cis-BP (20 Weirauch et al., 2014) and HOCOMOCO (10 Vorontsov et al., 2024) represented TF binding motifs derived from large-scale *in vivo* and *in vitro* studies.

A series of studies proposed the hierarchical classification of mammalian TFs based on their DNA-binding domains (DBDs) (TFClass, 21-24 Wingender, 2013; Wingender et al., 2013, 2015, 2018). Thus, hundreds of TFs representing nine specific superclasses and several dozens of the most abundant classes were defined. Recently, the TFClass framework has been applied to other eukaryotic taxa in the JASPAR database (19 Rauluseviciute et al., 2024); the focused study has detail the results for plants (Plant-TFClass, 25 Blanc-Mathieu et al., 2023). This advance promotes bioinformatics analysis in plants, as no new TF superclasses have been identified at the highest level of the hierarchy in plants compared to the nine currently known superclasses in mammals (24 Wingender et al., 2018). As previously expected (26 Riechmann et al., 2000), about half of all TF classes are common to mammals and plants.

However, enriched motifs identified by *de novo* motif searches do not necessarily imply TF binding specificity. Although peaks are generally assumed to represent genomic loci with motifs of some TFs, *de novo* motif discovery tools can identify not only such motifs but also motifs enriched across the genome as a whole. These motifs are unlikely biologically relevant, and hereinafter they are referred to as non-specific. The most known examples of non-specific motifs are polyA tracts. Such motifs refer to simple sequence repeats (SSRs), short tandem repeats with a monomer length of 1- 6 bp (27 Srivastava et al., 2019). All processing steps of ChIP-seq data analysis, including peak calling and *de novo* motif search require careful filtration of possible false positive non-specific motifs (28 Ross et al., 2013).

Long before the era of massive genome sequencing, the sizes of DNA sequence sets were very small and motif search tools took only one set of training sequences, putatively enriched with specific TFBS motifs (the generative learning principle, Lawrence et al., 1993). Later, the principle of discriminant learning was proposed in much more advanced tools designed to analyze whole genomes (30-32 Redhead and Bailey, 2007; Keilwagen et al., 2011; Simcha et al., 2012). This principle took two sets of sequences, the first one still implied specific TFBS motifs (foreground set, peaks), while the second (background set) had to neutralize false enrichment of non-specific motifs from the foreground set. Therefore, the choice of background sequences is a key step required to estimate the significance of motifs enrichment in peaks (32,33,11 Simcha et al., 2012; Boeva et al., 2016; Liu et al., 2018). Formally, the specific and non-specific motifs are correct results of *de novo* motif discovery.

Occasionally, due to missing or inadequate selection of background sequences, enrichment of non-specific motifs can compete with and even exceed enrichment for specific motifs. So far, many studies have promoted the concept of the synthetic sequences as efficient for evaluation of the performance of motif finding in ChIP-seq data. This concept implied the generation of synthetic sequences by Markov chains of various orders (33-36 Tompa et al., 2005; Boeva et al., 2016; Jayaram et al., 2016; Castellana et al., 2021), or these sequences were taken as a complete dictionary of k-mers, i.e. equal frequencies of nucleotides were presumed (6 Kulakovskiy and Makeev 2013). Markov modeling of expected word frequencies has been very popular since the k-th order Markov chain captures compositional biases represented at the level of all words of lengths from 1 to k+1. However, for different word lengths, spectra of word frequencies in different genomes showed behaviors, strikingly diverse from those expected in Markov modeling (37 Csurös et al., 2007). In accordance with this, a benchmarking analysis of distinct *de novo* motif finding tools demonstrated that the synthetic approach gave too optimistic estimates of performances (32 Simcha et al., 2012).

This review suggested that background sets consisting of genomic sequences were needed to rigorously test hypotheses. This option implies randomly chosen genomic regions with basic properties of sequences (such as length, nucleotide composition, location in the genome, etc.) that match those from the foreground sequences (31,38-41 Heinz et al., 2010; Keilwagen et al., 2011; Worsley Hunt et al., 2014; Dang et al., 2018; Tsukanov et al., 2022).

Although several tools allowed the generation or at least application of either genomic or synthetic background sequences (38,40,42-45 Sharov and Ko, 2009; Heinz et al., 2010; Dang et al., 2018; Khan et al., 2021; Bailey, 2021; Santana-Garcia et al., 2022), none of them recommended one option as superior to the other. For example, each of the popular *de novo* motif search tools such as Homer (38) and STREME (43) proposes only two options for generating background sequences. In addition to the option of a custom user-defined set of background sequences, the tools offer genomic and synthetic approaches, respectively.

Therefore, the discrepancy in the overall results of various popular *de novo* motif search tools is related not only to the peculiarities of their algorithms, but also to different options for selecting background sequences. The application of the most reasonable approach for generating background sequences by each tool will certainly improve the quality of its results. To date, only one study (39 Worsley Hunt et al., 2014) has attempted to compare systematically the synthetic and genomic approaches generating background sequences for subsequent *de novo* motif discovery. However, a small benchmark collection of 43 ChIP-seq datasets for only several TFs were considered. As mentioned above, last five years brought several specific databases focused on the uniform processing of ChIP-seq data (Cistrome DB, ReMap and GTRD; 12-14). Hence, a larger study comparing the results of different background sequence generation approaches is now possible. Nevertheless, no systematic large-scale studies comparing the sensitivity and specificity of applying genomic or synthetic background sequences for the *de novo* motif discovery in ChIP-seq data have been performed so far.

In the current study, we applied the benchmark collections of ChIP-seq data to compare the synthetic and genomic approaches generating the background sequences for *de novo* motif discovery. The ‘synthetic’ approach destroys the significant enrichment of any motifs through the shuffling of nucleotides preserving only the nucleotide composition. The ’genomic’ approach selects sequences from the reference genome or certain its part randomly, using for each peak its A/T nucleotide content, thereby modeling the expected content of non-specific motifs. We aimed to clarify in massive tests which of two approaches was more sensitive in detecting the known motifs of target TFs, and simultaneously was more specific in restriction of non-specific motifs of SSRs as possible false positives. We compiled the benchmark collections of ChIP-seq datasets for plants (*A. thaliana*) and mammals (*M. musculus* and *H. sapiens*). We generated the background sets with the genomic and synthetic approaches, and performed *de novo* motif discovery. We ranked the enriched motifs according to the significance of their enrichment. Then, we marked enriched motifs that were significantly similar to the known motifs of target TFs, as well as those for the motifs of SSRs. We concluded that the genomic approach, compared to the synthetic one, showed more reliable detection of the known motifs of target TFs and more rigorous exclusion of the SSR motifs as possible false positives.

Finally, we developed the AntiNoise command-line software package and its web service. They applied the genomic approach to extract background sequences for the foreground sets from ChIP-seq data of all most popular in massive analysis eukaryotic species from fungi to plants and mammals. The AntiNoise web service provides the opportunity for fast extraction of the sets of background sequences that in subsequent *de novo* motif search, through careful estimation of motif enrichment, potentiates deeper insight into yet hidden mechanisms of gene transcription regulation. The command-line software package has some additional options and it can be useful for the massive analysis.

## MATERIAL AND METHODS

### ChIP-seq data preparation

We extracted processed ChIP-seq data for *A. thaliana*, *M. musculus* and *H. sapiens* target TFs from GTRD (14). We selected ChIP-seq datasets that were preprocessed by the MACS2 peak caller (46) and had input control experiments in the primary processing pipeline. For *A. thaliana*, we extracted all available datasets, and for *M. musculus* and *H. sapiens*, we randomly selected about 15% of all available datasets from GTRD. The functionality of murine and human TFs were supported by their curated status (47 Lambert et al., 2018) (the high homology between human and murine TFs enabled this filtration, 23 Wingender et al., 2015), and for Arabidopsis TFs by annotations from PlantRegMap (48) and TAIR (49). For each ChIP-seq dataset, we defined 1000 top-scoring peaks as the foreground set for subsequent analysis. Each background set consisted of 5000 sequences, i.e. five background sequences for each foreground sequence were required. We applied ‘genomic’ and ‘synthetic’ approaches to generate for each foreground set the background sets of two types. The genomic approach chose sequences in the reference genome randomly: we allowed only the maximum deviation of 1% in the fraction of A/T nucleotides in each background sequence compared to the corresponding foreground one. The scheme in Figure 1 represents the algorithm of the genomic approach. Genomic sequence extraction requires either an unmasked or masked reference genome. A masked genome version can be prepared using the ’Exclusion of blacklisted regions’ or ’Retention of whitelisted regions’ options. The blacklisted option excludes certain genomic loci from the entire reference genome; the extraction procedure is applied to the remaining loci. The whitelisted option allows the extraction of background sequences only from certain specified regions; all other loci are excluded from the analysis. The synthetic approach performed the shuffling of nucleotides of peaks, exactly preserving a nucleotide content of each peak. Both approaches kept the lengths of sequences in the foreground and background sets unchanged.

**Figure 1.**
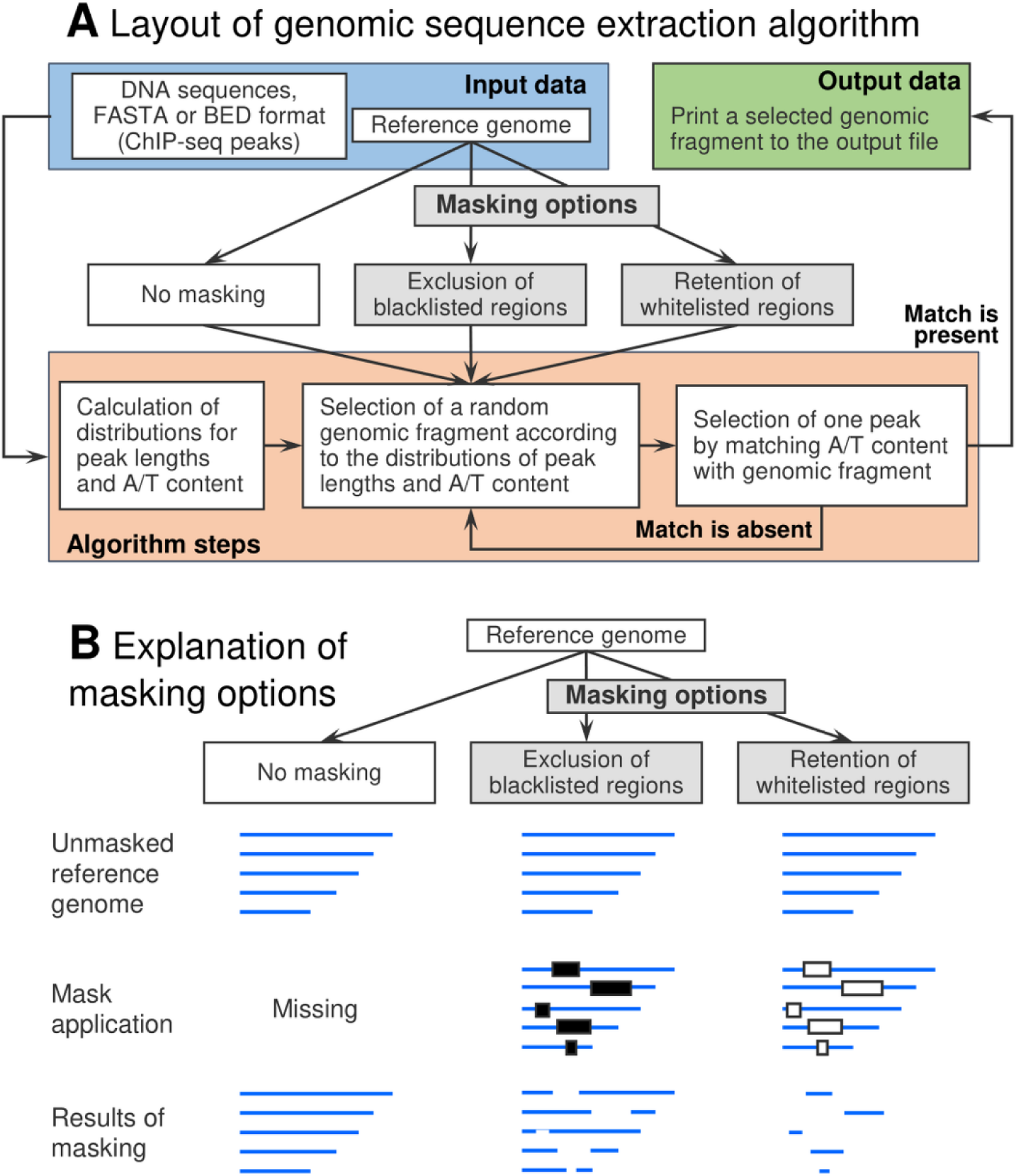
The algorithm of the genomic sequence extraction. (A) Layout of the algorithm. Blue color mark input data. First, input (foreground) sequences are used to compute the distributions of peak length and A/T content. Second, the reference genome is used to select proper background sequences. Hence, the output background sequences exactly match the foreground ones in length, while their A/T content allows only very small variation. Green color shows output data. (B) Scheme explaining two alternative masking options (grey). Here we either took the reference genome as is, the pathway ‘No masking’, or apply ‘Exclusion of blacklisted regions’, removing from the reference genome particular regions, and using all remaining loci in subsequent analysis, or apply another ‘Retention of whitelisted regions’, preserving in the reference genome certain specific regions and removing all remaining loci.

We required for each ChIP-seq dataset that a target TF respected the known motifs (as position frequency matrices) in its class/family from JASPAR (19 Rauluseviciute et al., 2024) or its family from Cis-BP (20 Weirauch et al., 2014). For this class or family, we required the presence of at least one member motif possessing the significant enrichment (p< 0.05, AME tool, 49) in the foreground set compared to the corresponding background set. Finally, murine/Arabidopsis benchmark ChIP-seq collections included 701/117 datasets for 212/57 target TFs, respectively (see Supplementary Tables S1, S2).

### ChIP-seq data analysis pipeline

For each pair of foreground and background sets we performed *de novo* motif discovery with the traditional motif model of a position weight matrix (PWM), STREME tool, (43).We used the default parameters of the STREME tool, the motif length varied from 8 to 15 nt. For each ChIP-seq dataset, the input data of this tool included a foreground set consisting of peaks and a background set generated from peaks using a synthetic or genomic approach; the output data were a ranking of the top ten enriched motifs according to the significance of their enrichment. We estimated the sensitivity and specificity of *de novo* motif discovery for all pairs of foreground/background sets and all enriched motif as follows. To estimate the sensitivity for a particular enriched motif, we tested its significance of similarity (p < 0.05) to all known motifs of TFs from the same class/family (JASPAR) or family (Cis-BP) as that of the target TF. To estimate the specificity for the same enriched motif, we tested whether the similarity of this enriched motif to any of the SSRs motifs was significant. We considered following SSRs motifs as possible false positives: mono-, di-, and trinucleotide repeats, i.e. two motifs of mononucleotide repeats A8 and G8, ten and 32 motifs of di- and trinucleotide repeats ((XY)4 and (XYZ)3, respectively). We used the TomTom tool (51) to assess the significance of similarity between motifs.

We applied Fisher exact test to estimate the significance of the difference between the numbers of datasets that had enriched motifs with certain ranks. First, Fisher test compared the number of datasets with enriched motifs with certain ranks between the genomic and synthetic approaches; these enriched motifs corresponded to motifs either of known TFs or SSRs (Table 1); we performed these tests separately for each collection. Second, Fisher test compared for a single approach the ranking of enriched motifs of target TFs or SSRs motifs in *A. thaliana* and one of mammalian (*M. musculus* or *H. sapiens*) benchmark collections (Table 2); we performed these tests separately for the genomic and synthetic approaches.

**Table 1.**
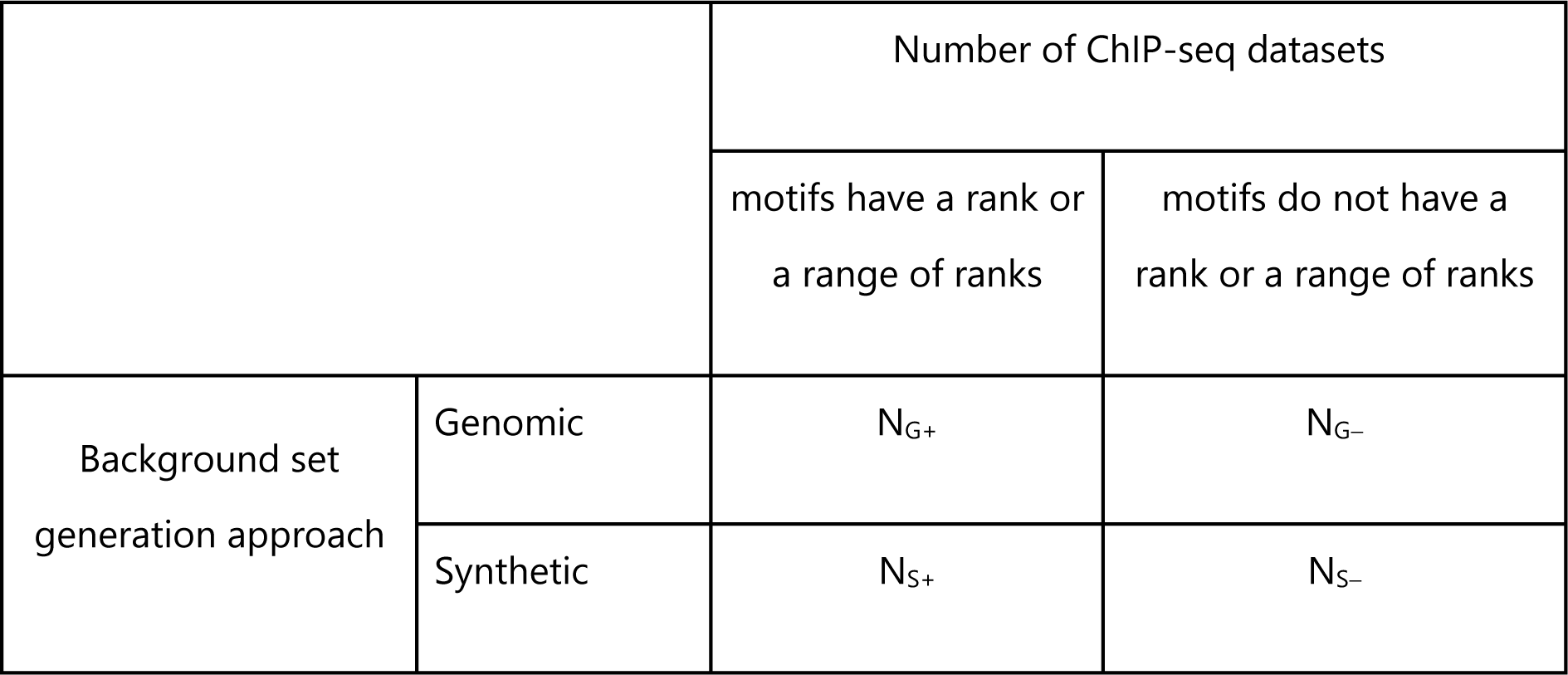
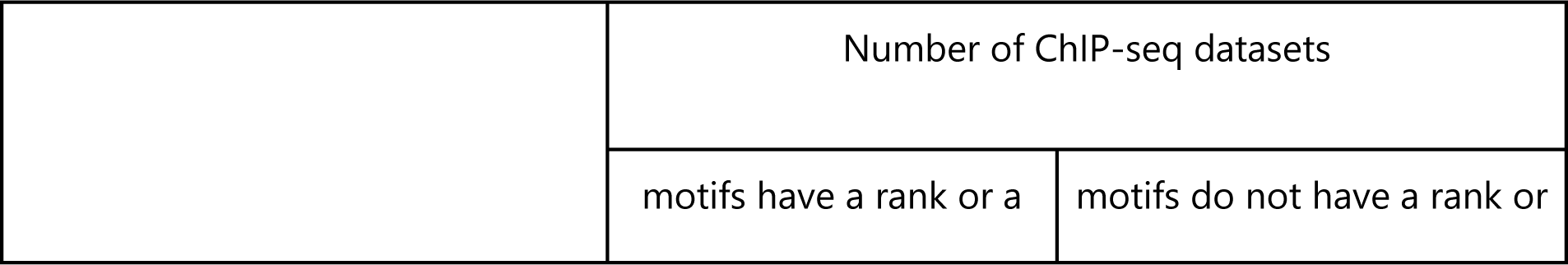
2 × 2 contingency table ‘Number of ChIP-seq datasets’ vs. ‘Background set generation approach’. The table counts ChIP-seq datasets with enriched motifs derived from *de novo* motif searches. These motifs have (+) or do not have (-) significant similarities to either known motifs of target TFs or SSR motifs. The table estimates the significant differences in ranking of enriched within the lists of top-ranked enriched motifs derived by *de novo* motif search for ChIP-seq datasets with application of the genomic and synthetic background sequences generation approaches. These tests were separately applied to the benchmark collections of *M. musculus* and *A. thaliana*.

**Table 2.** 2 × 2 contingency table ‘Number of ChIP-seq datasets’ vs. ‘Benchmark collection’. The table counts ChIP-seq datasets with certain enriched motifs derived from *de novo* motif search. These motifs have (+) or do not have (-) significant similarities to either known motifs of target TFs or SSR motifs. Table estimates the significant differences in ranking of enriched motifs within the lists of top-ranked enriched motifs derived by *de novo* motif search for ChIP-seq datasets from the benchmark collections of *M. musculus* (MM) and *A. thaliana* (AT). These tests were separately applied for the genomic and synthetic background sequences generation approaches.

|  |  | range of ranks | a range of ranks |
| --- | --- | --- | --- |
| Benchmark<br>collection | *M. musculus* | $N_{MM+}$ | $N_{MM-}$ |
| | *A. thaliana* | $N_{AT+}$ | $N_{AT-}$ |

We used the hierarchical classification of TFs by their DBDs (TFClass, 22-24; Plant-TFClass, 25) to study the relationship between the structure of DBDs of target TFs and efficacy of the genomic/synthetic background sequences application. For plant TFs we also used annotations from PlantRegMap (48) and JASPAR (19). Supplementary Tables S3 and S4 list superclasses and classes for all target TFs from two benchmark collections.

### Web service architecture

We proposed the web service AntiNoise promoting the genomic approach of background sequences extraction. The kernel of the web service was implemented in the C++ language. The kernel searched background sequences in the reference genome sequences (see above). The user interface (input/output data) and running of the kernel scripts were implemented in PHP language (version 7.4.3). In addition, Python code used matplotlib (52) and numpy (53) libraries to draw charts depicting distributions of the A/T content and dinucleotide frequencies.

## RESULTS

### Pipeline for ChIP-seq data analysis

We extracted ChIP-seq data for *M. musculus*, *H. sapiens* and *A. thaliana* target TFs from GTRD (14), we took each ChIP-seq dataset as the foreground set and generated for it background sets with the synthetic and genomic approaches (see Materials and Methods). Figure 1 explains the layout of the algorithm applied for the genomic approach of background sequences selection. For each set of foreground sequences, we computed the enrichment of the known motifs of target TFs respective to either the genomic or synthetic set of background sequences (AME tool, 50); this left in analysis 701/ and 117 ChIP-seq datasets for *M. musculus*, *H. sapiens* and *A. thaliana* (see Materials and methods). *De novo* motif discovery (STREME tool, 43) provided the top ten enriched motifs for each ChIP-seq dataset. Then, we estimated the significances of similarity between these enriched motifs and known motifs of TFs from the same class/family as a target TF. We applied at this step the motif library from JASPAR (19) and Cis-BP (20). Similarly, we estimated the significance of similarity between enriched motifs and the motifs of SSRs (see Materials and methods). Supplementary Tables S1 and S2 for all datasets provide ID of ChIP-seq datasets from GTRD, target TF names, the enrichment p-value (AME), the ranks of enriched motifs, the significances of motifs similarity, and the descriptions of the respective motifs from JASPAR/Cis-BP. Finally, we applied Fisher exact test to compare the sensitivity and specificity between the genomic and synthetic approaches, as well as the ranking of the motifs respecting the known motifs of target TFs or the motifs of SSRs, between the benchmark collections of ChIP-seq datasets for *M. musculus*/*H. sapiens* and *A. thaliana*.

### ChIP-seq data analysis pipeline

Figure 2A displays the analyses of two example ChIP-seq datasets for target TFs from *M. musculus* and *A. thaliana* (see row titles). Two columns depict the results for the genomic and synthetic approaches. Axes Y of four plots means the significances of the first ten enriched motifs from the results of *de novo* motif discovery as a function of their A/T content. For both examples from *M. musculus* and *A. thaliana*, the genomic approach assigned the first ranks to the known motifs of target TFs, but the synthetic approach displayed SSR motifs at the first ranks, so that motifs of target TFs had lower enrichments and subsequent ranks. These examples suggested that the genomic approach showed both better sensitivity and specificity than the synthetic approach.

**Figure 2.**
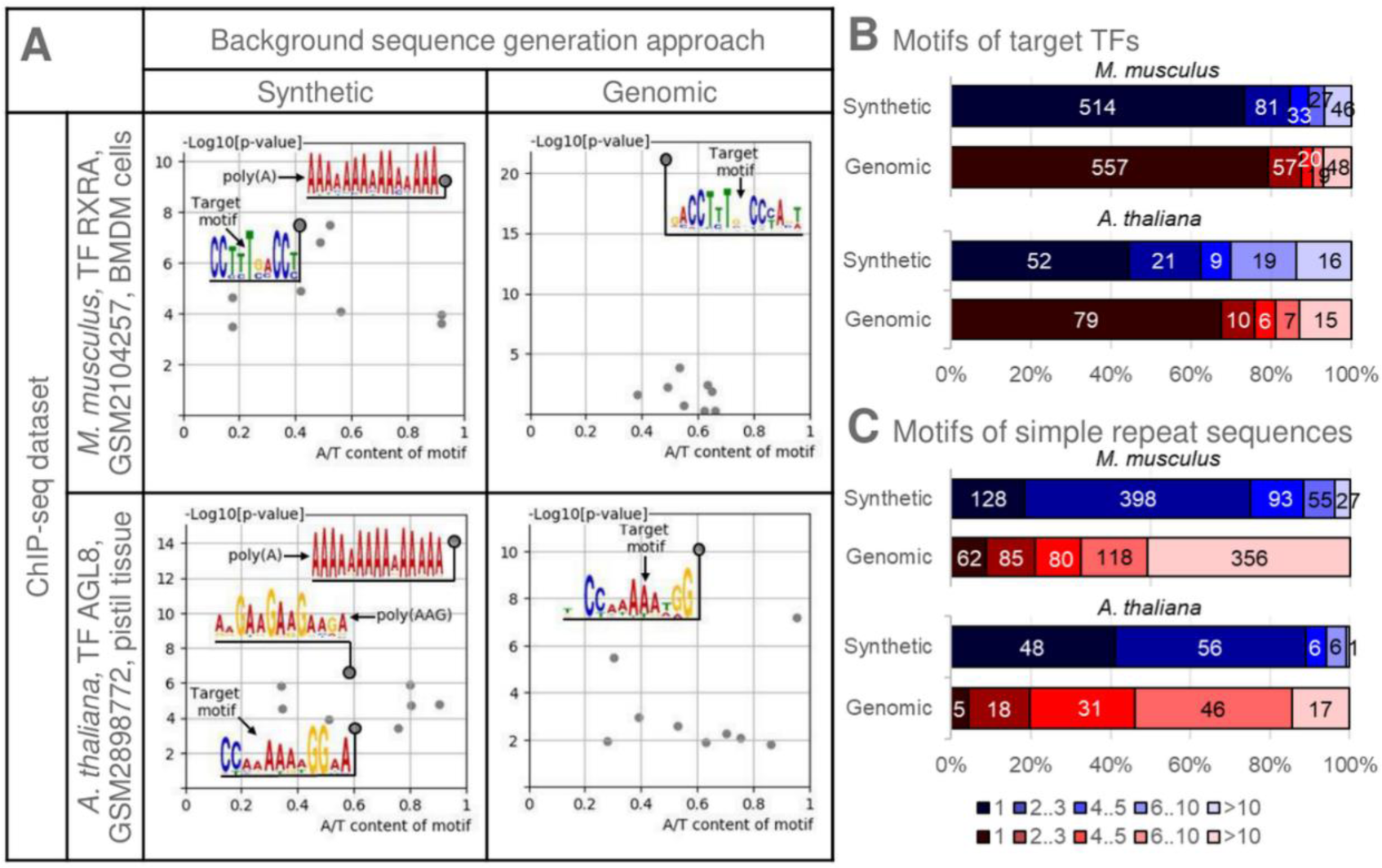
Abundances of motifs of target TFs and motifs of SSRs revealed by *de novo* motif discovery with application of the synthetic and genomic background approaches. (A) The ranking of motifs of target TFs and SSRs in the results of *de novo* motif discovery for two example ChIP-seq datasets. Columns show the results of *de novo* motif search for the synthetic and genomic background datasets. Rows represent the analysis of *M. musculus* and *A. thaliana* ChIP-seq datasets (TF RXRA, GTRD PEAKS039042, GEO GSM2104257, bone marrow-derived mesenchymal stem cells; TF AGL8, GTRD ID PEAKS042554, GEO GSM2898772, pistil tissue). Axes X and Y imply A/T content of motifs and the significance of motif enrichment from the STREME tool, –Log10[p-value]. This enrichment reflects the rank of a motif in the result of *de novo* motif search. Arrows and logos mark motifs of target TFs and motifs of SSRs. (B)/(C) The distributions of ranks of enriched motifs that are significantly similar to either the known motifs of target TFs (B) or SSRs (C). The distributions derived from the results of *de novo* motif discovery with application of the synthetic or genomic approaches for the benchmark collections of *M. musculus* and *A. thaliana* ChIP-seq data. Axes X mark the number of datasets possessing certain ranks of enriched motifs in the lists from *de novo* search; axes Y imply the synthetic and genomic background approaches. Red/blue colors mark the distributions computed for genomic/synthetic background sets; the darker/lighter shades of colors show the numbers of datasets possessing motifs with higher/lower ranks of enriched motifs.

Next, we analyzed two benchmark collections of ChIP-seq data. The distributions of ranks of the motifs of target TFs for two collections confirmed the conclusions derived from the analysis of two examples (Figure 2B). Thus, for *A. thaliana* and *M. musculus*, the genomic approach retained motifs significantly similar to known motifs of target TFs at the first ranks in 79 and 557 datasets, correspondingly, out of total 117 and 701 datasets. For the synthetic approach, the corresponding numbers were notably smaller (52 and 514). Respective distributions for SSR motifs showed the opposite trend (Figure 2C). The genomic approach revealed notably fewer datasets with SSR motifs at the first ranks (5/62 for *A. thaliana* / *M. musculus*) compared to the synthetic approach (48/128).

Application of Fisher exact test (Table 1) confirmed that the fractions of motifs corresponding to target TFs and ranked first were significantly higher for the genomic approach than for the synthetic approach (for the collections of *M. musculus* and *A. thaliana*, p < 0.01 and p < 0.001, see Figure 3A). The subsequent ranks showed the same trend, although it gradually became insignificant. The first-ranked enriched SSR motifs occurred significantly less frequently in lists derived from the genomic approach than from the synthetic approach (p < 5E-7 and p < 5E-12, Figure 3B). The subsequent ranks showed the similar and sometimes even more significant depletion.

**Figure 3.**
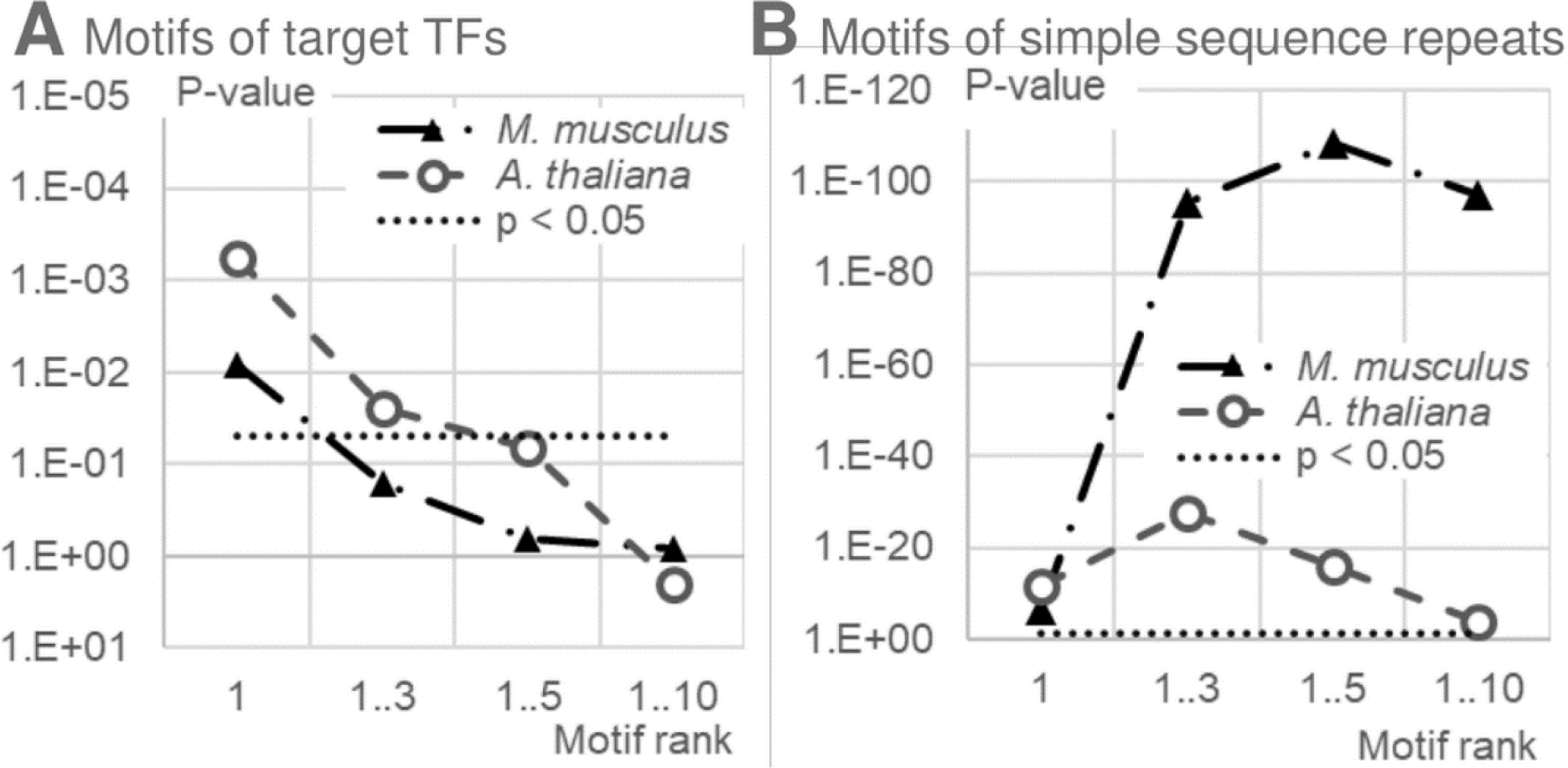
The significance of difference in ranking of the enriched motifs between the synthetic and genomic background approaches. *De novo* motif search revealed the ranks of enriched motifs. The Fisher exact test separately for the *M. musculus* / *A. thaliana* collections compared the fractions of datasets possessing enriched motifs with certain ranks. Panels (A)/(B) display the analysis for the enriched motifs, significantly similar to the motifs of target TFs and the motifs of SSRs. Axes X and Y show the motif rank and the significance by exact Fisher test, respectively; dotted lines mean the significance threshold (p < 0.05). Table 1 explains the Fisher task applied in analysis.

Next, we applied Fisher exact test separately for each of the background generation approaches to compare *M. musculus* and *A. thaliana* collections (Table 2). We revealed that the fractions of datasets possessing the enriched motifs respecting target TFs are significantly higher for the *A. thaliana* collection compared to those for the *M. musculus* collection. In this comparison, the genomic and synthetic approaches achieved moderate and high significance, p < 1E-2 and p < 1E-8 (Figure 4A). The comparison between the fractions of datasets containing enriched SSR motifs (Figure 4B) showed that they were significantly more abundant in *A. thaliana* than in *M. musculus*. This trend was significant for the top ranks of enriched motifs (1 and 1-3) for the synthetic approach, and for the wider ranges of ranks (1-5 and 1-10) for the genomic approach. Thus, for the synthetic approach, the enriched motifs with first ranks were SSRs much more frequently in the *A. thaliana* ChIP-seq data collection than in the *M. musculus* collection (p < 2E-7). Notably, the genomic approach rejected the significance of the corresponding trend for the enriched SSR motifs of the first ranks.

**Figure 4.**
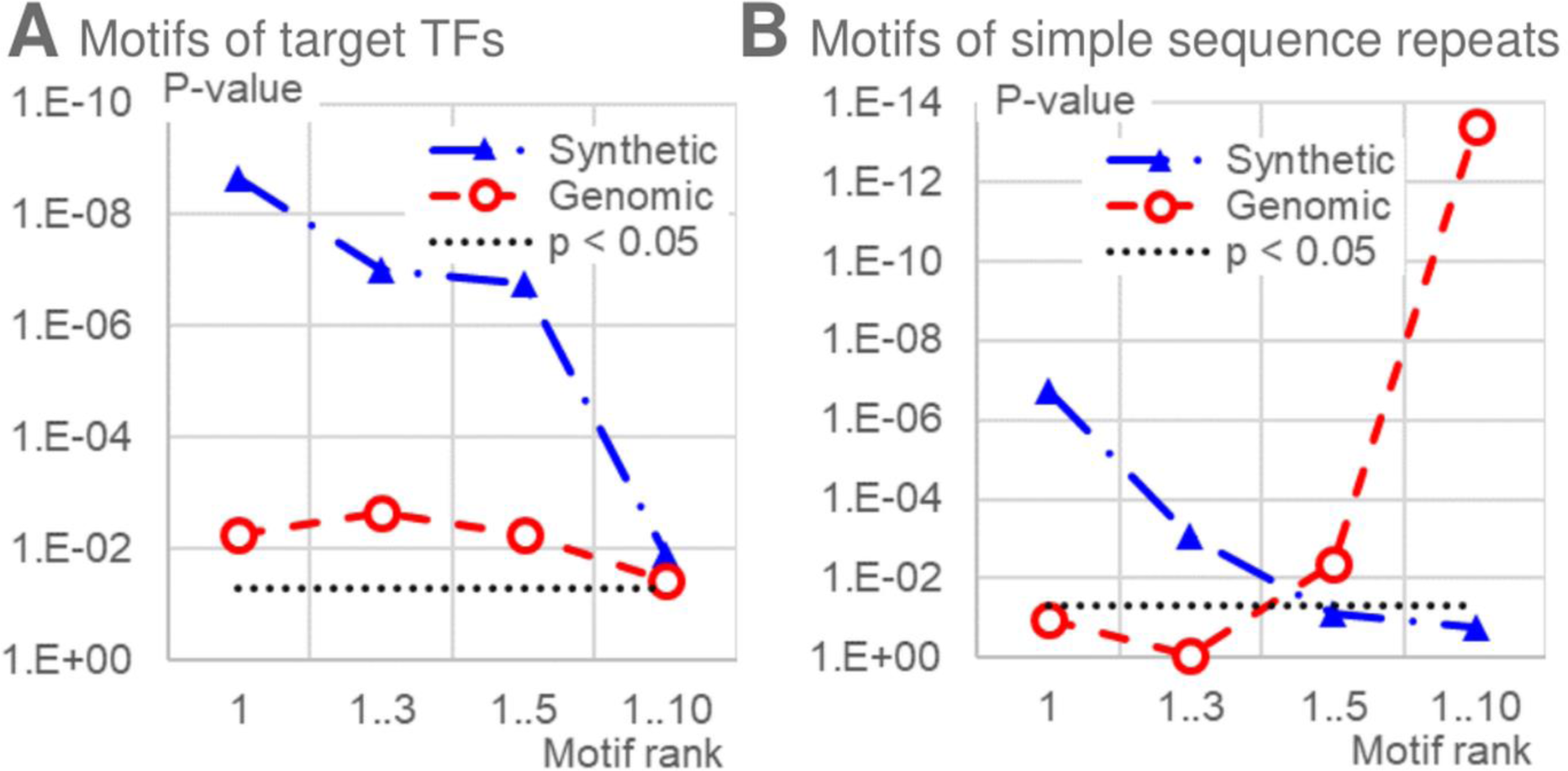
The significance of difference in ranking of the enriched motifs between the *M. musculus* and *A. thaliana* benchmark collections. *De novo* motif search revealed the ranks of enriched motifs. The Fisher exact test separately for the synthetic and genomic background approaches compared the fractions of datasets possessing enriched motifs with certain ranks. Panels (A)/(B) display the analysis for the enriched motifs, significantly similar to the motifs of target TFs and the motifs of SSRs. Axes X and Y show the motif rank and the significance by Fisher exact test, respectively; dotted lines mean the significance threshold (p < 0.05). Table 2 explains the Fisher task applied in analysis.

Overall, we found that the genomic approach of background sequences generation provided both better sensitivity and specificity for benchmark ChIP-seq data collections from mammals and plants. As expected, we confirmed that the synthetic and genomic approaches guaranteed the significant enrichment of motifs for target TFs in *de novo* motif discovery results for all ChIP-seq data collections. In contrast to the genomic approach, the synthetic approach assigned the highest ranks to the enriched SSR motifs substantially more frequently in the *A. thaliana* ChIP-seq data collection compared to the *M. musculus*. Hence, *de novo* motif discovery in plant ChIP-seq data requires more careful processing taking into account possible false positives of SSR motifs enrichment.

### Genomic approach shows better sensitivity for target TFs of almost all classes

Next, we tested whether the difference between the results of applying the genomic and synthetic approaches depended on the structure of DBDs of target TFs. We used annotations of *M. musculus* and *A. thaliana* TFs from JASPAR database, and applied their hierarchical classification into the superclasses and classes (see Materials and methods, Supplementary Tables S3 and S4, 21-25). Figure 5 for the most abundant superclasses and classes of target TFs of *M. musculus* and *A. thaliana* shows the distributions of the number of datasets, with enriched motifs, that are significantly similar to known motifs of target TFs with certain ranks in the lists of motifs obtained using the genomic and synthetic approaches. Supplementary Tables S5 and S6 present the results of corresponding analyses for all classes. Overall, for almost all the most abundant classes of *M. musculus* and *A. thaliana* target TFs (12 of 13), the genomic approach is more sensitive than the synthetic approach. Only for the class of C2H2 zinc finger factors in *M. musculus* synthetic/genomic approaches shows the first ranks for 124/122 datasets out of 137. Since these numbers are almost equal, one exception still confirmed the general trend.

**Figure 5.**
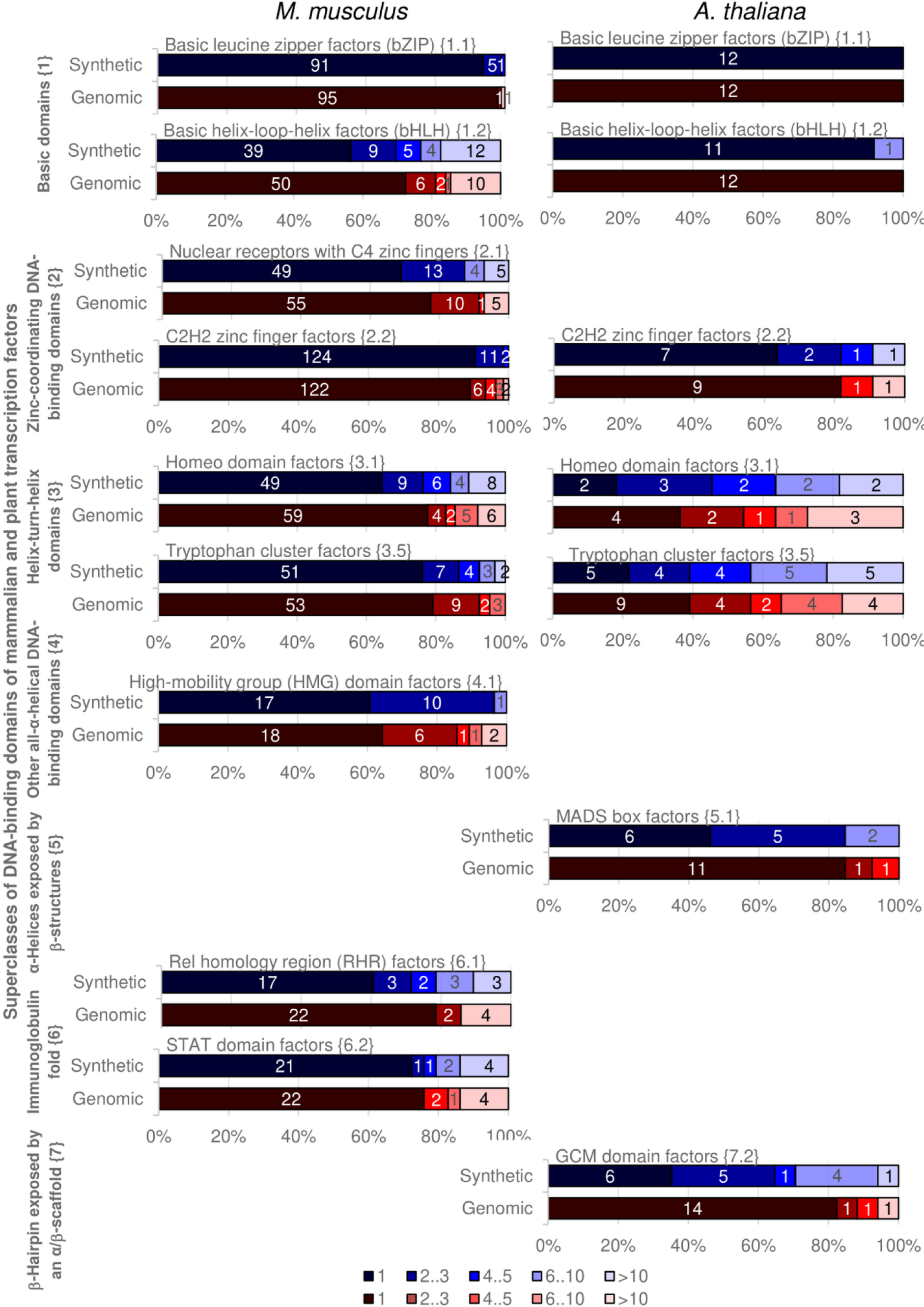
Application of the genomic and synthetic approaches for target TFs of various DBD structures. Left/right columns present analysis of ChIP-seq datasets from the *M. musculus* and *A. thaliana* collections. The hierarchical classifications of *M. musculus* and *A. thaliana* target TFs by the structure of DBDs were derived from TFclass and Plant-TFclass (22–25), see Materials and methods. Axes X mark the number of datasets possessing certain ranks of enriched motifs in the lists from *de novo* search; axes Y imply the application of the synthetic and genomic background approaches for ChIP-seq data with target TFs from various classes. Green arrows and numbers in square brackets mean the names of TF superclasses (Basic domains {1}, Zinc-coordinating DNA-binding domains {2}, Helix-turn-helix domains {3}, Other all-α-helical DNA-binding domains {4}, α-Helices exposed by β-structures {5}, Immunoglobulin fold {6}, β-Hairpin exposed by an α/β-scaffold {7}); the diagrams are entitled with the names of TF classes. The ranks for TFs were assigned by the enrichment significance of motifs from *de novo* motif discovery for ChIP-seq datasets for the *M. musculus* and *A. thaliana* benchmark collections. Only the most abundant classes were represented. See Supplementary Tables S3 / S4 for the respective ranking of enriched motifs for all TF classes in *M. musculus* / *A. thaliana*.

### Anti-Noise web service and command line software package: input/output data and functionality

For a wider application of the genomic approach in further researches, we implemented it in the AntiNoise web service as the command line software package and web service. The command line software package is available at https://github.com/parthian-sterlet/antinoise. The package allows application of genomic and synthetic approaches. Additionally, the package provides perl scripts starting from the reference genome sequences in one FASTA file. This file contains all chromosomes, it can be found in any genome-specific database. There are three scripts defined by source of background genomic sequences:

- ‘No masking’. The entire reference genome. In this case, no genome masking is performed.
- ‘Blacklisted region masking’. The entire reference genome with certain excluded ‘blacklisted’ regions. We mask these regions, thereby they completely excluded from output data. We propose ‘blacklisted’ regions from a recent study for mouse and human as examples (54).
- ‘Retention of whitelisted regions’. The background sequences are restricted to genomic ‘whitelisted’ regions, and all remaining genomic loci were masked. We propose the promoter regions of all protein-coding genes (-5000; +100) as default option of the whitelisted regions for all species.

Since regions in the same list, no matter blacklisted or whitelisted regions, can overlap each other, we took this into account and provided an internal step that clears possible self-overlap for a script that handles any of these masking options.

The web service is available at https://denovosea.icgbio.ru/antinoise/. Figure 6A displays the main and advanced options of the web service:

- Genome release and species, among them the animals human (Homo sapiens, hg38), mouse (Mus musculus, mm10), rat (Rattus norvegicus, Rnor_6.0), zebrafish (Danio rerio, GRCz11), fly (Drosophila melanogaster, dm6) and roundworm (Caenorhabditis elegans, WBcel235); the plants are arabidopsis (*A. thaliana* TAIR10), soybean (Glycine max v2.1), maize (Zea mays, B73), and liverwort (Marchantia polymorpha, MpTak v6.1); the fungi baker’s yeast (Saccharomyces cerevisiae, R64-1-1) and fission yeast (Schizosaccharomyces pombe, ASM294v2);
- Required number RBF of background sequences per one foreground sequence. The default value RBF = 5 implies that that if calculations are successively completed for all NF input sequences, then totally NB = RBF * NF background sequences are found;
- Either ‘no masking’, or ‘Retention of whitelisted regions’ options are applied. The ‘whitelisted regions’ are the promoter regions of all protein-coding genes, (-5000; +100) relative to the 5’ ends of genes.
- Deviation δ of the A/T nucleotide content of each background sequence from that for the corresponding foreground sequence. The default value 0.01 allows the mismatch of one bp per a sequence length of 100 bp in a foreground sequence.
- Threshold FMIN for the minimum fraction of completely processed foreground sequences to stop calculations. The default value of 0.99 means that calculations stop if for 99% of all foreground sequences for each sequence at least RBF background sequences are found for each sequences;
- Maximal number of attempts NA to find matching background sequences in the genome. If a given number NA of last attempts to find any at least one more background sequence are unsuccessful, the algorithm terminates. The default value 50000.

**Figure 6.**
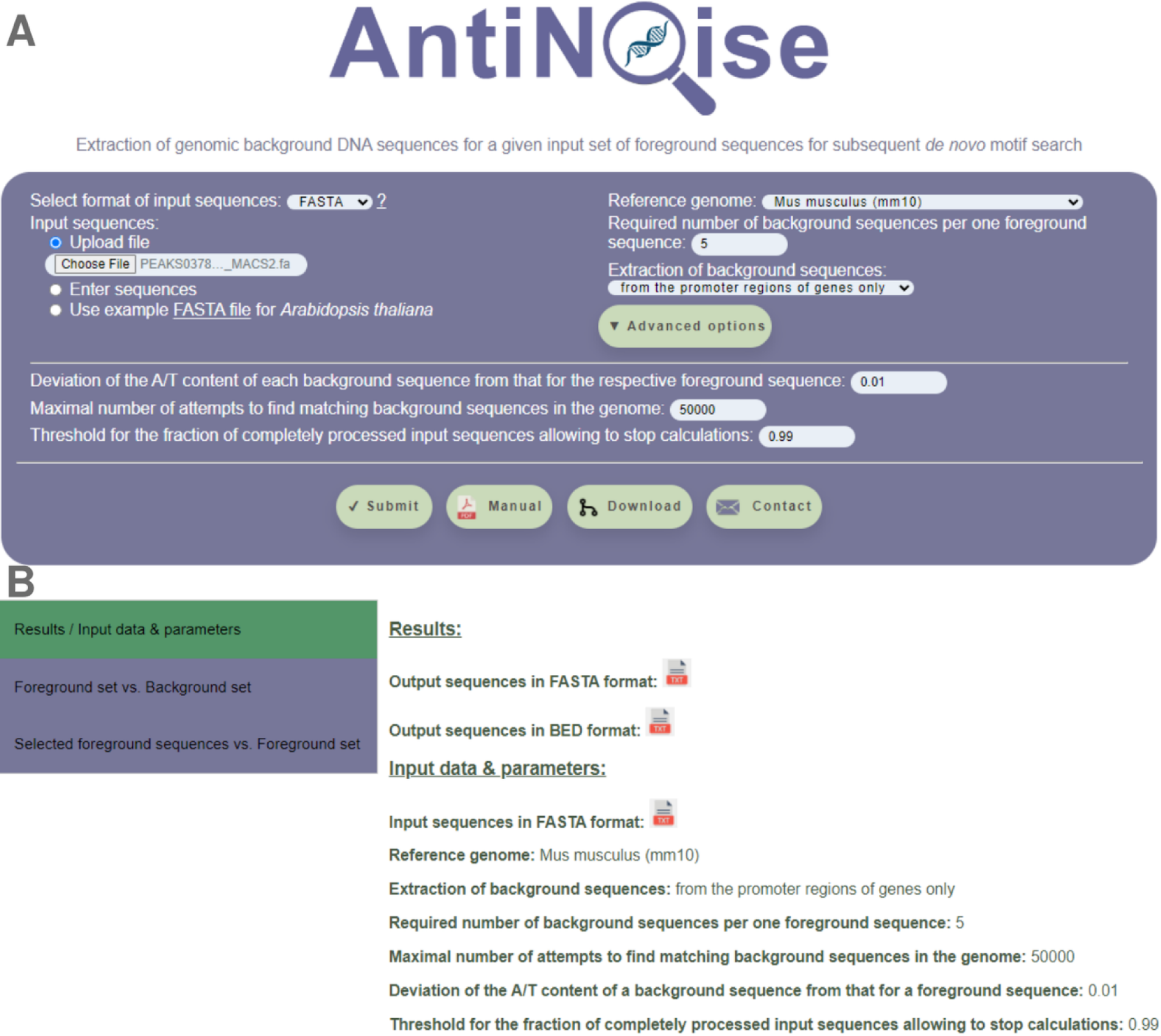
Main screenshots of the AntiNoise web application. (A) Application page with an example. Here user can enter input sequences, set the parameters of a calculation task and start calculations. (B) The ‘Results / Input parameters’ output tab provides access to input (foreground) sequences and output (background) sequences, and lists major options of a calculation task. ChIP-seq dataset for *M. musculus* TF MEF2A, GEO GSM1629389, GTRD ID PEAKS037873 is used as an example.

Figure 6A shows the screenshot for the application page of the web service with an example. Here we analyzed the dataset of 1000 top-scored ChIP-seq peaks for *M. musculus* cortical neurons treated with 10nM purified recombinant Reelin for 1 hour, TF MEF2C, GEO GSM1629389 (55), GTRD ID PEAKS037873.

Pressing the ’SUBMIT’ button starts the calculation and provides a web link to the page with its results. During the running process, the service indicates the number of found background sequences. The output data of the web service include a link to a file with the main calculation result, a set of genomic background sequences in FASTA format. These results respect the first tab ‘Results / Input parameters’ (Figure 6B). In addition, four charts on two tabs illustrate the validity of the background sequences search. The second tab ‘Foreground set vs. Background set’ shows two charts depicting the distributions of the A/T content and of the dinucleotide frequencies for the foreground and background sets (Figure 7A). These charts can be critical for choosing between various types of background sets. The third tab ‘Selected foreground sequences vs. Foreground set’ demonstrate two charts listing foreground sequences that did not reach the required number of background sequences per one foreground sequence. These two charts apply the metrics of the A/T content and dinucleotide frequencies to compare the selected foreground sequence with the whole set of foreground sequences (Figure 7B). While the A/T content means the metrics applied for the genomic background sequences selection, the dinucleotide frequencies as the k-mers of the shortest length 2 bp show the behavior of the simplest motifs. These two charts detect foreground sequences with abnormal mono- and dinucleotide content.

**Figure 7.**
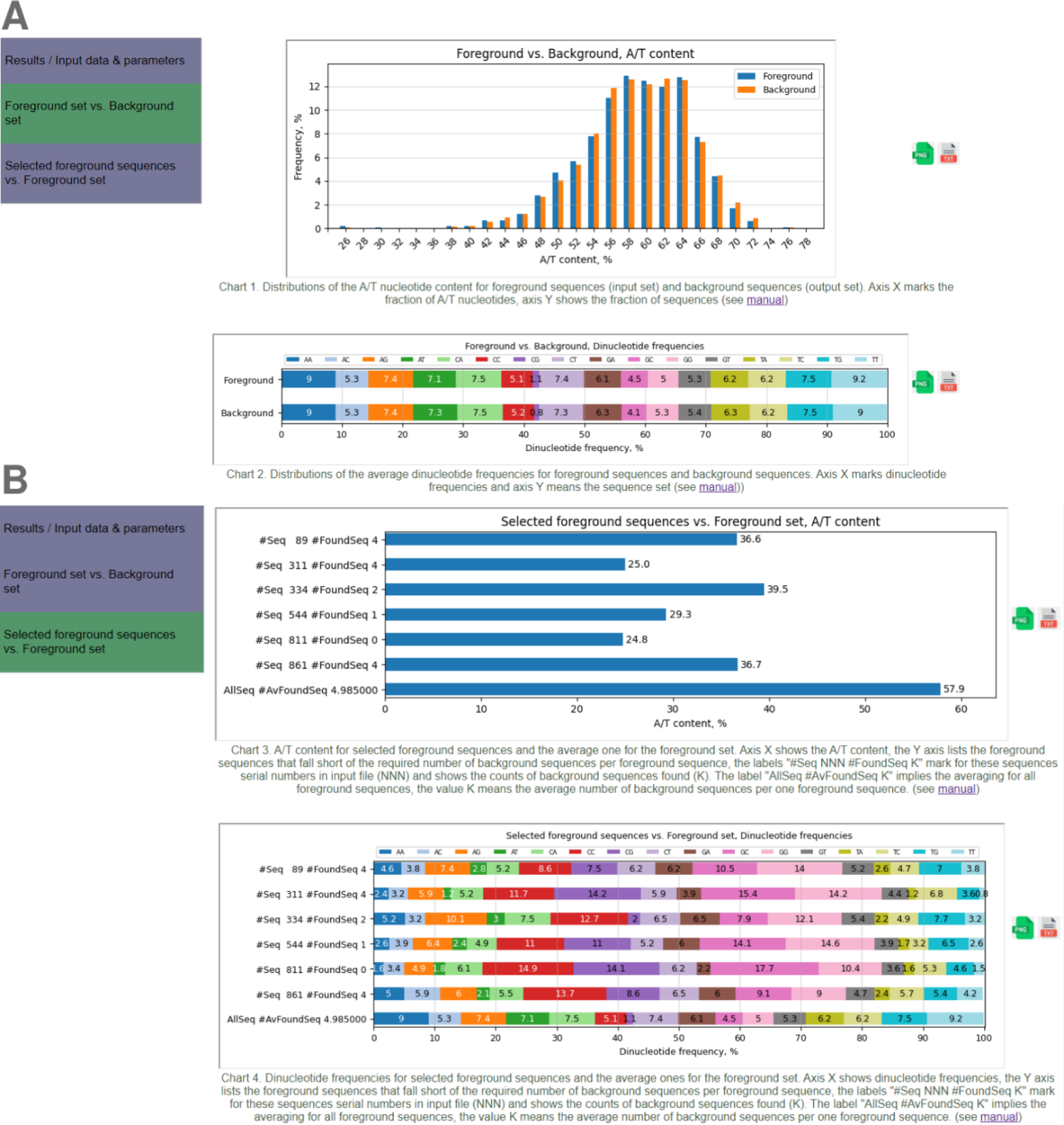
Additional output screenshots of the AntiNoise web application. (A) The ‘Foreground set vs. Background set’ output tab compares the input and output sequences. One chart depicts the A/T content and another shows dinucleotide frequencies. (B) The ‘Selected background sequences vs. Foreground’ output tab compares input sequences that did not reach the required threshold of found genomic background sequences and all input sequences. the Y axis lists foreground sequences that did not reach the required number of background sequences per one foreground sequence, the labels “#Seq NNN #FoundSeq K” mark for these sequences serial numbers in input file (NNN) and shows the counts of background sequences found (K). The label “AllSeq #AvFoundSeq K” implies the averaging for all foreground sequences, the value K means the average number of background sequences per one foreground sequence. Axes X show the A/T-content (top) and dinucleotide frequencies (bottom). The ChIP-seq dataset for *M. musculus* TF MEF2A, GEO GSM1629389, GTRD ID PEAKS037873 is used as an example.

## DISCUSSION

In the current study, we considered the potency of two popular approaches to generating background sequences for subsequent *de novo* motif search in ChIP-seq data. The synthetic approach performs nucleotides shuffling that abolishes the enrichment of any motifs. This procedure radically destroys in the foreground sequences the enrichment of k-mers of any length. These k-mers represent either specific or non-specific motifs; they compete between each other at the next step of *de novo* motifs search. Applying the synthetic background approach inevitably results in a lower frequency of non-specific motifs in the background sequence compared to the background sequence. In this case, the enrichment of non-specific motifs in the foreground set is not due to binding of specific TFs, but rather corresponds to the overall specificity of the full-length genome in terms of oligonucleotide composition. Attempting to suppress these noisy genomic bias motifs in the foreground set, one should try to preserve their content in the background set, while destroying the enrichment of context-specific motifs presumed to have TF binding functionality. Hence, the genomic approach implies the extraction of the background sequences from the reference genome sequences.

Before the beginning of the next-generation sequencing era, both TFBS and genomic mapping data were scarce, and, conventionally, motif discovery algorithms applied the synthetic sequences to model the expected frequencies of motifs (34 Tompa et al., 2005). However, application of these algorithms for newly available large datasets derived from ChIP-seq technology (56 Johnson et al., 2007) raised many unanswered issues, including long computation time, too redundant output data, variation in a threshold for the peak quality, filtration of genome bias motifs, etc. (6,9,28,32 Simcha et al., 2012; Zambelli et al., 2013; Kulakovskiy and Makeev 2013; Ross et al., 2013). In addition, the chromatin as primary source of ChIP-seq data often complicated the detection of motifs of target TFs due to their possible indirect binding through intermediate proteins, co-binding with partner TFs, and structural heterogeneity within the same TFs (57,58 Chumpitaz-Diaz et al., 2021; Yu et al., 2021). Thus, even a variety of distinct algorithmic strategies gave only a limited success in higher eukaryotes (3,9,36,43,59 Zambelli et al., 2013; Tran and Huang, 2014; Lai et al. 2019; Bailey, 2021; Castellana et al., 2021).

Earlier, it was found that the distributions of the relative abundance of short oligonucleotides were strikingly diverse among DNA sequences from the genomes of various eukaryotic taxa, and these distributions were certainly different from those expected from Markov modeling (60 Karlin and Ladunga, 1994). Namely, the di-/tri-nucleotide frequencies in genomic sequences differed markedly from those expected by their mono-/di-nucleotide content, etc.

Authors explained these differences through distinct structural properties of short k-mers, in particular, the base-step stacking capacities, duplex curvature and other higher order DNA structural features of dinucleotides (61 Karlin and Burge, 1995). It was concluded that the genome-wide consistency of dinucleotide relative abundance values suggested involvement of the fundamental biological processes, such as DNA replication, recombination and repair. Later, the analysis of the whole chromosomes of various species from distant eukaryotic taxa confirmed that the deviations of dinucleotide frequencies from those expected according to their nucleotide content were distinctly genome-specific (62 Gentles and Karlin, 2001). These studies showed that the sequence bias in whole genomes implied the species-specific pattern of enriched and depleted oligonucleotides of various lengths, including lengths typical for TFBS motifs (6-20 bp). The synthetic approach applies Markov models with various orders m of a chain, which may be ranged from 0 to 5 (63,64 Siebert and Söding, 2016; Eggeling et al., 2017). Therefore, even the largest order of a Markov chain can ensure the preservation of k-mers of short lengths, up to hexamers. However, the occurrence in the foreground sequences of k-mers of longer lengths, particularly those with lengths as long as the TFBS motifs (6-20 bp), can be wrongly regarded as enrichment. Hence, the synthetic approach can create an artificial enrichment of non-specific motifs in the foreground set.

Thus, because the genomic approach reflects the genome-specific bias in oligonucleotide frequencies it should be superior to the synthetic approach. The synthetic approach has been very popular in motif discovery tools to generate the background sequences (9,65,66 Jiang et al., 2008; Zambelli et al., 2013; Bailey et al., 2011). Some modern tools do not allow to adopt the genomic background sequences for *de novo* motif discovery (64,67-69 Kulakovskiy et al., 2010; Keilwagen and Grau, 2015; Eggeling et al, 2017; Caldonazzo Garbelini et al, 2018), while others allow both synthetic or genomic background sequences (38,43,45,70 Heinz et al., 2010; Kiesel et al., 2018; Bailey, 2021; Santana-Garcia et al., 2022). Alternative suggestion for *de novo* motif discovery compiled genome sequences flanking peaks to the background set (7172Guo et al., 2011; Samee et al., 2019). However, even the peak calling tool selection and its options influence on the precise positioning of peaks borders. E.g., the peak callers GEM/MACS2 provide peaks of fixed/varied lengths (71,46) (Guo et al., 2011; Zhang et al., 2008). Therefore, we suspect that the compilation of background sequences from areas flanking the peaks is not a completely correct methodology. Among the current sources of TFBS motifs derived from ChIP-seq data, three are the most reliable and popular, the HOCOMOCO (10), CisBP (20) and JASPAR (19). Among them, only the CIS-BP was developed with the support from the various types of genomic background sequences (73 Weirauch et al., 2013).

In the current study, we took in analysis ChIP-seq data from the GTRD, since this database combined uniformly processed chromatin immunoprecipitation data with detailed annotations, such as the application of the input control experiment in the processing pipeline, descriptions of tissue/cell/treatment conditions, application of various peak caller tools, etc. We retained for analysis only ChIP-seq datasets with enrichment (AME) of the known motifs of TFs from the same classes/families (JASPAR) or families (Cis-BP) as the target TFs. Thus, popular motif databases supported the context specificity of target TFs binding. We treated enriched SSR motifs as potential false positives. For each ChIP-seq dataset, we considered the top ten most enriched motifs from results of *de novo* motif discovery (STREME). This choice was due to the inability of ChIP-seq technology to distinguish between direct and indirect TF-DNA interactions (1,2,59 Nakato and Shirahige, 2017, Lai et al., 2019; Lloyd and Bao, 2019), and target TFs could have lower ranks in the output lists. In confirmation of this, many massive applications of *de novo* motif discovery to ChIP-seq data showed that about a half of datasets did not reveal the known motifs for target TFs as the first-ranked enriched motifs (41,74-76 Worsley Hunt and Wasserman, 2014; Levitsky et al., 2019; Tsukanov et al., 2022; Karimzadeh and Hoffman, 2022). A rank of each enriched motif allowed estimating the sensitivity and specificity of each of the background generation approaches, through estimates of the significance of the similarity of that enriched to either known motifs of target TFs or SSR motifs, respectively.

To select sequences into the background set, we applied two metrics of genomic DNA loci. The first is A/T content, the variation of which also accurately reflects the variation of G/C content, reflecting the ratio of the content of relatively strong G/C and weak A/T pairs of nucleotides with three and two hydrogen bonds in DNA. For example, it is difficult to find any G/C-reach motifs of the KLF/SP1 family in A/T-rich genomic loci. Accurate retention of the second metric, the sequence length, is required to support popular measures of accuracy for *de novo* motif search, such as the Precision-Recall curve or Area Under Curve (ENCODE-DREAM *in vivo* TF Binding Site Prediction Challenge, (77)).

Example analysis of two ChIP-seq datasets (Figure 2A) showed that the use of genomic background sequences compared to synthetic ones provided higher enrichment of known motifs of target TFs and lower enrichment of SSR motifs. We proposed that these differences were due to the complete destruction of the enrichment of any motifs by nucleotide shuffling as the generation procedure of synthetic sequences, while the procedure of the genomic approach accounted the expected content of genome bias motifs in ChIP-seq data.

The systematic analysis of the *M. musculus*, *H. sapiens* and *A. thaliana* benchmark collections of ChIP-seq data confirmed these conclusions (Figure 2B,C). We came to two concordant conclusions. First, known motifs of target TFs showed significantly higher ranks in the results of *de novo* motif discovery for the genomic approach compared to those for the synthetic approach (Figure 3A). Second, SSR motifs demonstrated significantly lower ranks in the results of *de novo* motif discovery for the genomic approach compared to those for the synthetic approach (Figure 3B).

Next, we compared the results of separate applications of either genomic or synthetic approach between ChIP-seq data collections for *M. musculus* and *A. thaliana*. Both collections revealed the significant enrichment of known motifs of target TFs for the *A. thaliana* collection, although the synthetic approach showed higher significance (Figure 4A). Surprisingly, the synthetic approach provided substantially higher enrichment of SSR motifs for the *A. thaliana* collection compared to the *M. musculus* collection (motif rank 1, 1-3, Figure 4B). The genomic approach did not show the significant enrichment for the same ranks of the enriched motifs. Hence, whereas both approaches are sensitive due to enrichment of the known motifs of target TFs, the specificity appreciated through the abundances of potentially false positive motifs of SSRs is substantially worse for *A. thaliana* ChIP-seq data over those for *M. musculus*.

Finally, we considered the hierarchical classification of target TFs from ChIP-seq data of *M. musculus* and *A. thaliana* by their structure of DBDs (22–25,19). We showed that the higher enrichment of the known motifs of target TFs in the results from the genomic approach compared to those from the synthetic one is observed for almost all most abundant classes of murine or Arabidopsis TFs (Figure 5).

A necessary step of the processing of massive sequencing data such as ChIP-seq is the assessment of motif enrichment reflecting the binding specificity of target TFs. The choice of a particular approach has been essential for both specific tools generating background sequences (44,65 Jiang et al., 2008; Khan et al., 2021), and for *de novo* motif search tools (38,67,78 Heinz et al., 2009; Kulakovskiy et al., 2010; Thomas-Chollier et al., 2012), and for the special databases of TFBS motifs (19,20). However, until now, no one has performed a massive analysis of ChIP-seq data for various eukaryotic taxa to compare two the most popular approaches - the generation of background sequences by the synthetic or genomic approaches. Since our study resulted in very strong arguments in favor of the genomic approach (Figures 2-5), we implemented it as the command line software package and the web service (Figures 6,7). They allow extract background sequences for the most popular in massive sequencing analysis eukaryotic genomes from yeasts to mammals and plants. Thus, we propose a flexible approach to robustly support the identification of specific TF targeting motifs in widely used *de novo* motif search tools from massive sequencing data.

A recent comprehensive all-against-all TF binding motif benchmarking study (79 Ambrosini et al., 2020) showed that TF binding specificity correlates with the structural class of its DBD. Thus, different PWMs for TFs from the same structural classes tended to perform similarly across experiments; and the best performing PWM model often respected the same TF class. Another conclusion from the benchmarking study (79 Ambrosini et al., 2020) stated that the practical set of motifs for many biological applications is much smaller than the number of motifs already contained in the most popular motifs collections. Therefore, we hope that the main conclusions of our study concerning the advantages of the genomic background sequences over synthetic sequences hold true for target TFs from a variety of eukaryotic taxa and for target TFs of any structural class.

We performed a *de novo* motif search for benchmark collections of ChIP-seq data for target TFs from mouse, human and Arabidopsis. We aimed to investigate whether background sets consisting of genomic or synthetic sequences provide both a higher prediction rate of known motifs of target TFs and a lower prediction rate of potentially false positive SSR motifs. We found that although both genomic and synthetic approaches provide pronounced enrichment of known target TF motifs, the synthetic approach compared to the genomic approach yields a very significant increase in the proportion of SSR motifs representing possible false positives. We confirmed the advantage of the genomic approach over the synthetic approach in terms of sensitivity of detection of motifs of known target TFs for almost all the most common classes of target TFs in mammals and plants. As for specificity, the use of the synthetic approach compared to the genomic approach more often resulted in higher ranks of enriched SSR motifs for plant than for mammalian ChIP-seq data. To summarize, massive analysis of mammalian and plant ChIP-seq data has shown that the genomic approach is more effective than the synthetic approach in generating background sequences for *de novo* motif discovery. Therefore, we propose to use genomic sequences extracting as a default option for generating a set of background sequences when applying *de novo* motif search to ChIP-seq data. To promote and widely apply the results of our analysis, we implemented the genomic approach of background sequences generation as the web service as the AntiNoise web service and provided its extended options in the command line version.

## Supporting information

Graphical Abstract

## DATA AVAILABILITY

### SUPPLEMENTARY DATA

Supplementary Data are available online.

### AUTHOR CONTRIBUTIONS

V.V.R.: Formal analysis, Visualization. A.V.T. Formal analysis. A.G.B.: Software, Writing—review & editing. V.G.L.: Conceptualization, Investigation, Methodology, Validation, Software, Supervision, Validation, Visualization, Writing—original draft, Writing—review & editing.

## ACKNOWLEDGEMENTS

The bioinformatics data analysis was performed in part on the equipment of the Bioinformatics Shared Access Center within the framework of State Assignment Kurchatov Genomic Center of ICG SB RAS [FWNR-2022-0020].

