## Supplementary figures and images for "Genomic background sequences systematically outperform synthetic ones in de novo motif discovery for ChIP-seq data"

### Graphical Abstract

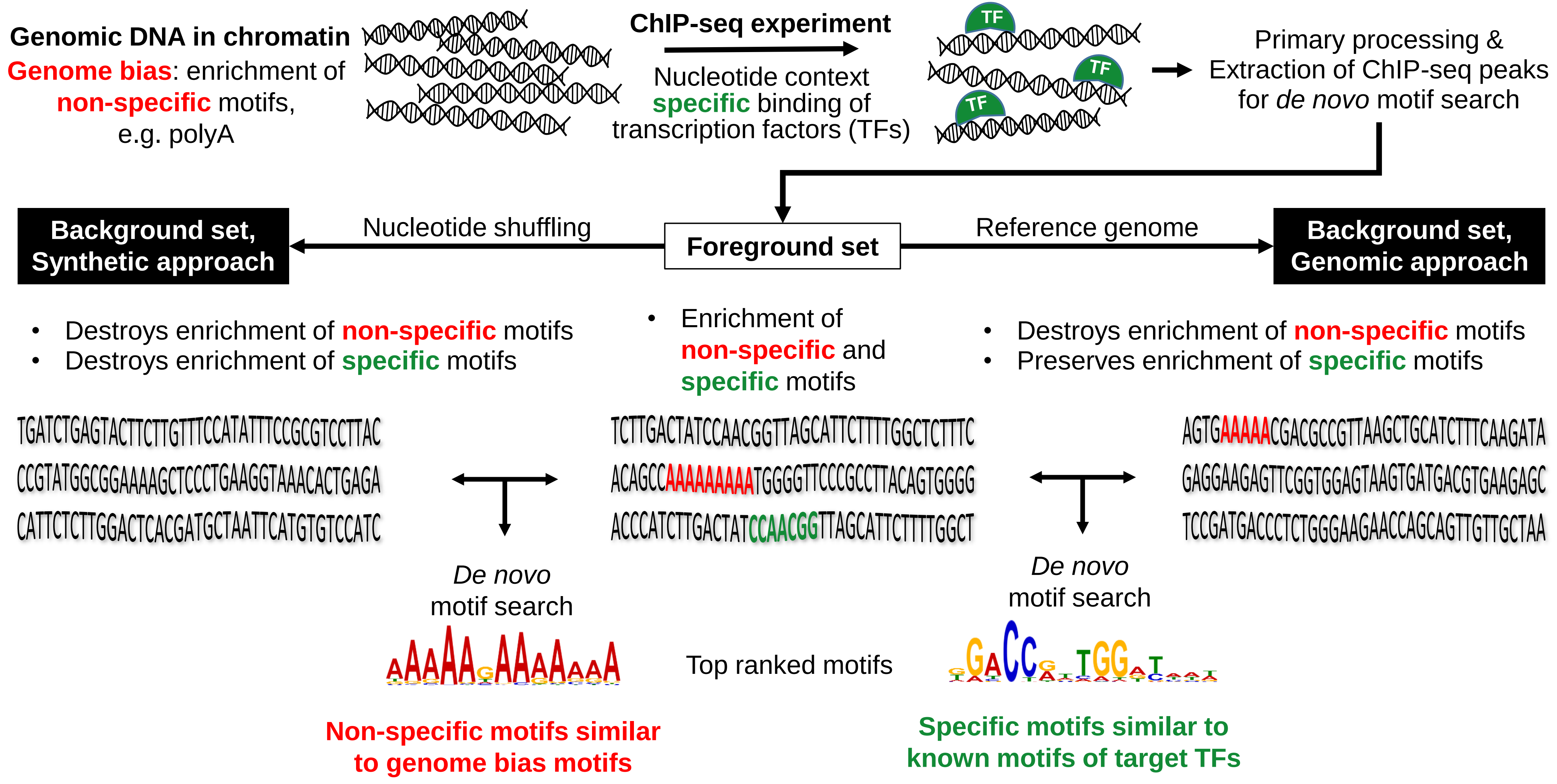
